# Structured light projection using image guide fibers for in situ photo-biofabrication

**DOI:** 10.1101/2024.12.10.627729

**Authors:** Parth Chansoria, Michael Winkelbauer, Shipin Zhang, Jakub Janiak, Hao Liu, Marcy Zenobi-Wong

## Abstract

Light-based biofabrication techniques have revolutionized the field of tissue engineering and regenerative medicine. Specifically, the projection of structured light, where the spatial distribution of light is controlled at both macro- and micro-scale, has enabled precise fabrication of complex three-dimensional structures with high resolution and speed. However, despite almost two decades of progress, biofabrication processes have been mostly limited to benchtop devices which limit the flexibility in terms of where the fabrication can occur. Here, we demonstrate a Fiber-assisted Structured Light (FaSt-Light) projection apparatus for rapid in situ crosslinking of photoresins. This approach uses image-guide fiber bundles which can project bespoke images at multiple wavelengths, enabling flexibility and spatial control of different photoinitiation systems and crosslinking chemistries and also the location of fabrication. We demonstrate coupling of different sizes of fibers and different lenses attached to the fibers to be able to project small (several mm) or large (several cm) images for material crosslinking. FaSt-Light allows control over the cross-section of the crosslinked resins and enables the introduction of microfilaments which can further guide cellular infiltration, differentiation and anisotropic matrix production. The proposed approach could lead to a new range of in situ biofabrication techniques which improve the translational potential of photo-fabricated tissues and grafts.

## Introduction

Past two decades have seen a tremendous growth in the field of light-based biofabrication.^[1,2]^ A variety of processes have emerged to crosslink biomaterials,^[3]^ or to change functional properties of the materials across spatial domains.^[2]^ These processes offer high precision (up to 150 nm with two-photon polymerization (2PP)),^[4]^ speeds (printing in several seconds using volumetric bioprinting),^[5,6]^ and scalability (complex constructs spanning several centimetres can be printed using digital light projection (DLP)).^[7]^ These different approaches have found applications in the fabrication of acellular grafts for tissue repair,^[8,9]^ or for biofabrication using living cells to engineer a variety of tissues such as cartilage,^[10,11]^ liver,^[12,13]^ cardiac^[14,15]^ and skeletal muscle,^[16,17]^ etc. Spatially controlled light projections have also been used for inducing functional changes in photo-responsive materials, such as spatially-tuned presentation of growth factors or molecules for cell differentiation.^[18–20]^

While the light-based techniques highlighted above can be used to fabricate complex constructs for a variety of regenerative applications, these approaches have been mostly limited to benchtop devices.^[21,22]^ The constructs are fabricated over a substrate or within a printing vial, and later post-treated (washing, secondary crosslinking, additional sterilization, etc.), and cultured (for constructs with cells) before being used for implantation.^[21,22]^ A benchtop fabrication approach limits the region and the containers these materials could be printed into. Furthermore, for applications where the constructs are being used for implantation, handling and surgical implantation of the grafts is often complicated. Notably, several studies have demonstrated in situ photo-crosslinking of hydrogels for applications including diabetic wound healing,^[23,24]^ treatment of myocardial infarction,^[25,26]^ or corneal regeneration,^[27,28]^ etc. However, these approaches have crosslinked bulk resins around the tissue by projecting homogeneous beams through fibers or handheld light sources. Here, controlling the micro- and macro-architecture of the in situ crosslinkable photoresins (usually hydrogels for biofabrication applications) can allow better cellular infiltration, growth and tissue formation. To print complex structures, approaches such as tomographic printing would be complicated for in situ printing as it requires light projection from multiple resin orientations.^[21]^ In contrast, 2PP or DLP techniques have been used, since the light is projected from one direction.^[29–31]^ For instance, Urciolo and colleagues demonstrated 2PP *in vivo* bioprinting using a near infrared (NIR) laser (850 nm), which could penetrate up to several millimeters under the skin.^[32]^ In this study, a laser beam was moved to trace the outline of the constructs layer-by-layer, to create arbitrary shapes subcutaneously. In a similar study, Chen and colleagues demonstrated subcutaneous printing of photoresins by projecting images at NIR wavelength (980 nm),^[30]^ where crosslinking was achieved using up-conversion nanoparticles coated with the photoinitiator lithium phenyl-2,4,6-trimethylbenzoylphosphinate (LAP). In contrast to these approaches, we hypothesized that coupling the projection apparatus to image guide fiber bundles could allow flexibility in terms of guiding the light to harder-to-reach places (e.g., tissues inside the body or in different types of resin containers, etc.). Image guide fiber bundles, also called coherent image guides or fiber optic imaging bundles, are stacked bundles of optical fibers which can convey optical spatial properties from one end of the fiber to another.^[33,34]^ These fibers have been extensively used for imaging in regions where the approach area is small and difficult to reach (e.g., endoscopy).^[35,36]^ The captured image is guided through the bundles into a camera sensor, which can then discern the depth and colour of the images and render it on a user-interface.^[33,34]^ Interestingly, reversing the function of the image guide fibers, where custom images generated from a projection apparatus are guided through the fibers for the crosslinking of materials, has not yet been explored for biofabrication applications. Notably, such a system is different from recent approaches using an optical fiber which emits a uniform light beam, and where the fiber needs to be mechanistically traversed to create complex crosslinked constructs.^[37]^

In this work, we demonstrate a **F**iber-**a**ssisted **St**ructured **Light** (**FaSt-Light**) projection apparatus (illustrated in **Figure 1**) for rapid in situ photo-biofabrication using image guide fibers. The aspect of structured light entails two scales of control over the spatial distribution of light: **1.** Macroscale structures (> 50 µm) imparted through the digital micromirror device (DMD) in the projection system which controls the image at multiple wavelengths projected through the fibers, allowing control over the cross-section of the crosslinked resin, and **2.** Microscale control, which refers to the micro-filamentation of light beams due to the optical modulation instability when the laser speckles interact with the photoresin, allowing the introduction of cell-guiding microfilaments (2-8 µm) within the crosslinked resin. We have previously leveraged the combined effect of macro- and micro-scale structures for the fabrication of microfilamented scaffolds using filamented light (FLight) biofabrication, which is a benchtop approach to generate highly aligned tissue constructs.^[10,16]^ In this work, we demonstrate a new system, which generates speckled images at multiple light wavelengths (405, 450, 520 nm), which are projected through image guide fiber bundles onto photoresins contained within in a variety of enclosures (cuvettes, well plates, or defects on the tissues). We investigate the resins crosslinked using the FaSt-Light approach in terms of the resulting macro and micro-scale features and further elucidate the effects of fiber cross-section area, projection distance, and lenses coupled to the fibers on the structures. Through *in vitro* experiments, we show that the FaSt-Light apparatus can be used to biofabricate viable tissue constructs or grafts which promote cell-guidance and infiltration. Furthermore, we also demonstrate selected proof-of-concept applications where the approach could be used for the crosslinking of structured photoresins *in vivo*, which can lead to new approaches for tissue regeneration.

**Figure 1.**
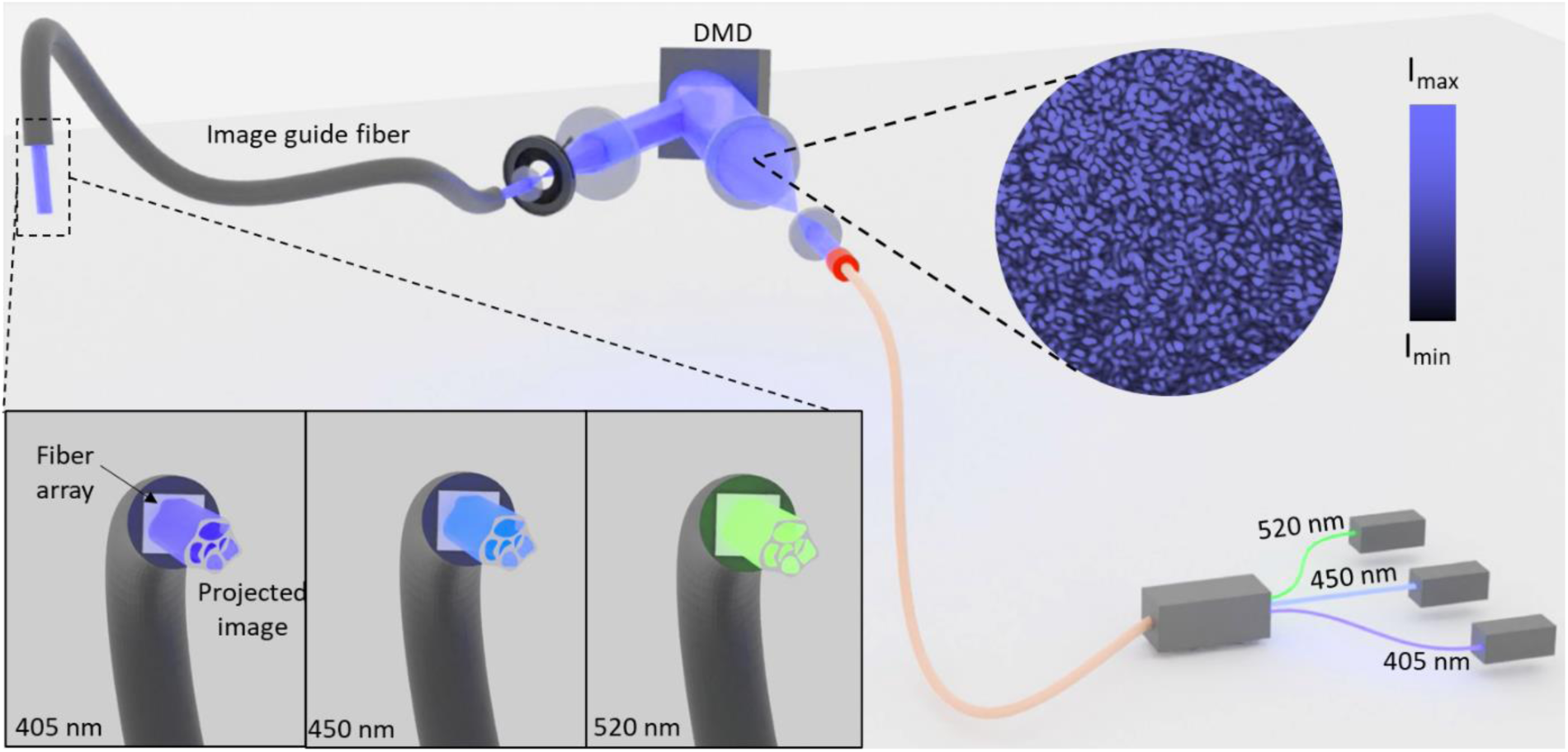
Schematic of the FaSt-Light apparatus capable of projecting bespoke image patterns of different wavelengths through an image guide fiber bundle. Note that the contrast ratio of the speckle patterns (intrinsic to the laser light engine) has been increased in the illustration to demonstrate the effect of light filamentation due to optical modulation instability.

## Results

The assembly of the FaSt-Light apparatus is illustrated in **Figure 1**. The projection system in the apparatus comprises of a light engine (i.e., laser modules), a digital micromirror array device (DMD) and a set of telescopic lenses before and after the DMD. The light engine for the three laser modules (continuous wave emission at 405, 450 and 520 nm) is connected to a single output fiber, which allows homogenization of the laser light beam at each wavelength and a consistent projection into the first telescopic lens set. The first telescopic lens expands the light to be able to cover the entire mirror array in the DMD. The image is then shaped using the DMD and then travels through the second set of telescopic lenses with an iris in-between the lens to remove the auxiliary images arising from the DMD. The projected image is then directed through the image guide fiber bundle, coupled to the projection setup, into the photoresin.

For the photocrosslinking schemes used in this work, we used 5% w/v gelatin methacryloyl (GelMA) in phosphate buffered saline (PBS) as the base material, for it is one of the most widely-used photoresin in light-based biofabrication.^[38–40]^ We further used three different photoinitiation mechanisms which have absorption maxima near each wavelength. For photocrosslinking at 405 nm, we used the Norrish Type-I photoinitiator Lithium phenyl-2,4,6- trimethylbenzoylphosphinate (LAP), which undergoes a homolytic cleavage upon light absorption, producing free radicals which lead to covalent bonding of the methacrylate groups in the resin.^[41]^ For 450 nm, we used a redox-based photoinitiation system containing ruthenium(II) complex (Ru) and sodium persulfate (SPS).^[41]^ Here upon photo-absorption, the excited ruthenium(II) complex (Ru(bpy)₃²⁺) transfers an electron to sodium persulfate (SPS), generating sulfate radicals which initiate the photocrosslinking process. Finally, for crosslinking at 520 nm, we used a Norrish Type-II initiation system consisting of Eosin Y (EY), Triethanolamine (TEOA) and N-vinylpyrrolidone (NVP).^[41]^ In this system, photoabsorption at 520 nm leads to excitation of EY, followed by electron abstraction from TEOA to create amine radicals which initiate the crosslinking process. Here, NVP acts as a chain transfer agent and a reactive diluent, accelerating the crosslinking process and enhancing the overall polymerization efficiency.

We first conducted tests on the projection system alone (illustrated in **Figure 2A**) without the image guide fiber bundle, to determine the light doses for crosslinking of GelMA at each wavelength. The resulting prints of ETH logo using the projection system at each wavelength (optimal light doses shown in Figure S1) and the concentration of the photoinitiators used therein are shown in **Figure 2B**. The resulting constructs made at each wavelength featured microfilaments generated by optical modulation instability due to the optical autocatalysis (i.e., self-focusing) of the speckled laser light beam within the photoresin.^[16,21]^ The stitched micrograph of the ETH logo (**Figure 2B**) shows the sectional view of the microfilaments, while a three-dimensional (3D) micrograph at each wavelength is shown in **Figure 2C**. While the photoinitiation mechanisms and the light doses for polymerization (Figure S1) were different for each wavelength, we did not observe differences in the microfilament diameter (typically 2-6 µm, **Figure 2D**), which can be attributed to the use of a single fiber optic cable connected to the output of the light engine. We also projected spoke wheel patterns (**Figure 2C**) within cuvettes to determine the minimum attainable feature size when crosslinked at each wavelength. Here, while the photoresins containing LAP (i.e., 405 nm) and EY-NVP-TEOA (i.e., 520 nm) demonstrated similar resolution (150-200 µm), the resins containing Ru-SPS (i.e., 450 nm) showed larger minimum feature sizes (200-310 µm). This reduced resolution in the Ru-SPS crosslinked GelMA could be attributable to the strong oxidation potential of the sulfate radicals in the crosslinking process, which can lead to the formation of tyrosyl radicals on the tyrosine residues within the gelatin backbone. These tyrosyl residues can form covalent dityrosine bonds through radical coupling, leading to increased non-specific crosslinking in the matrix, thereby affecting the resolution. While the above three photoinitiation strategies have been shown to be cytocompatible,^[21,41]^ we conducted additional viability tests with our system. For these experiments, we encapsulated myoblasts (C2C12; 10^6^ cells/ml) in separate formulations of GelMA with the three different photoinitiation systems and biofabricated cylindrical constructs by projecting circular images (ɸ = 750 µm) into 2 mm path length cuvettes at each wavelength. The viability of the cells at day three was greater 90% in each condition (**Figure 2E**), thereby demonstrating the cytocompatibility of the light doses used in the projection apparatus.

**Figure 2.**
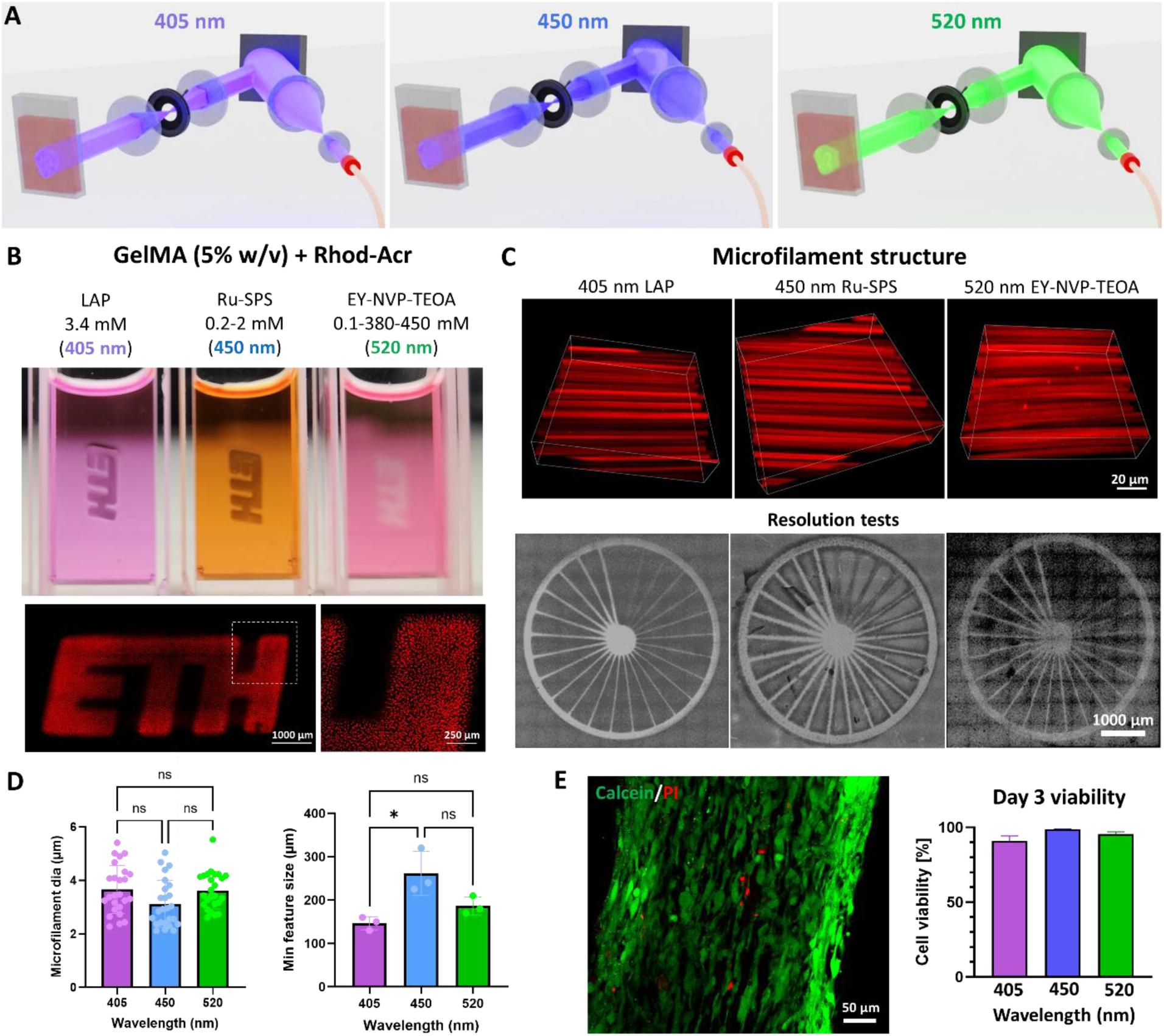
Validation of printing through the multi-wavelength structured light projection system. **A.** Illustration of laser activation to derive multiple wavelength projections (405, 450, 520 nm) into a photoresin cuvette. **B.** Crosslinked GelMA under various photoinitiation systems which have absorption maxima in the different wavelengths. The ETH logo demonstrates microfilaments generated due to optical modulation instability arising from the interaction of the light with the resin. **C.** Micrographs of the filament structures within the crosslinked constructs and the printed constructs for resolution tests. **D.** Results of microfilament diameter witihin the crosslinked constructs and minimum achievable feature size. * represents p < 0.05. **E.** Representative image of the viability of C2C12 cells (live cells are stained green with calcein AM and dead cells are stained red through propidium iodide; image shown is for the LAP photoinitiation system (405 nm)) at Day 3 within the crosslinked constructs and analysis of viability at each wavelength (i.e., for each photoinitiation system).

For subsequent studies we used the FaSt-Light apparatus, in which the image guide fiber bundle was coupled to the multi-wavelength projection setup. **Figure 3A** shows two different fiber bundles – one with thick cladding and one without - each featuring a 3×3 mm^2^ projection cross-section. Here, a thick cladding can offer enhanced protection and handleability to the fiber (e.g., resin crosslinking onto open tissue sites), while a thin cladding offers smaller form factor (e.g., suitability for endoscopy applications). Each of these fiber bundles feature stacked blocks of fiber arrays, where each block is an 8×8 grid of individual fibers. Here the arrangement of the blocks at the input of the fiber bundle matches that at the output, which is necessary for conveyance of the image features through the fiber bundle. Notably, the diameter of individual fibers (ɸ = 7.5 µm) in the fiber bundle is larger than pitch of the micromirror array (5.5 µm) in our projection setup, which limits the pixel size of the image projected through the fiber bundle to 7.5 µm. **Figure 3B** shows images (FaSt-Light and ETH logo) at each wavelength being projected using the two different fiber bundles (also see Video S1 and S2 of the image projection at each wavelength from the two fiber bundles).

**Figure 3.**
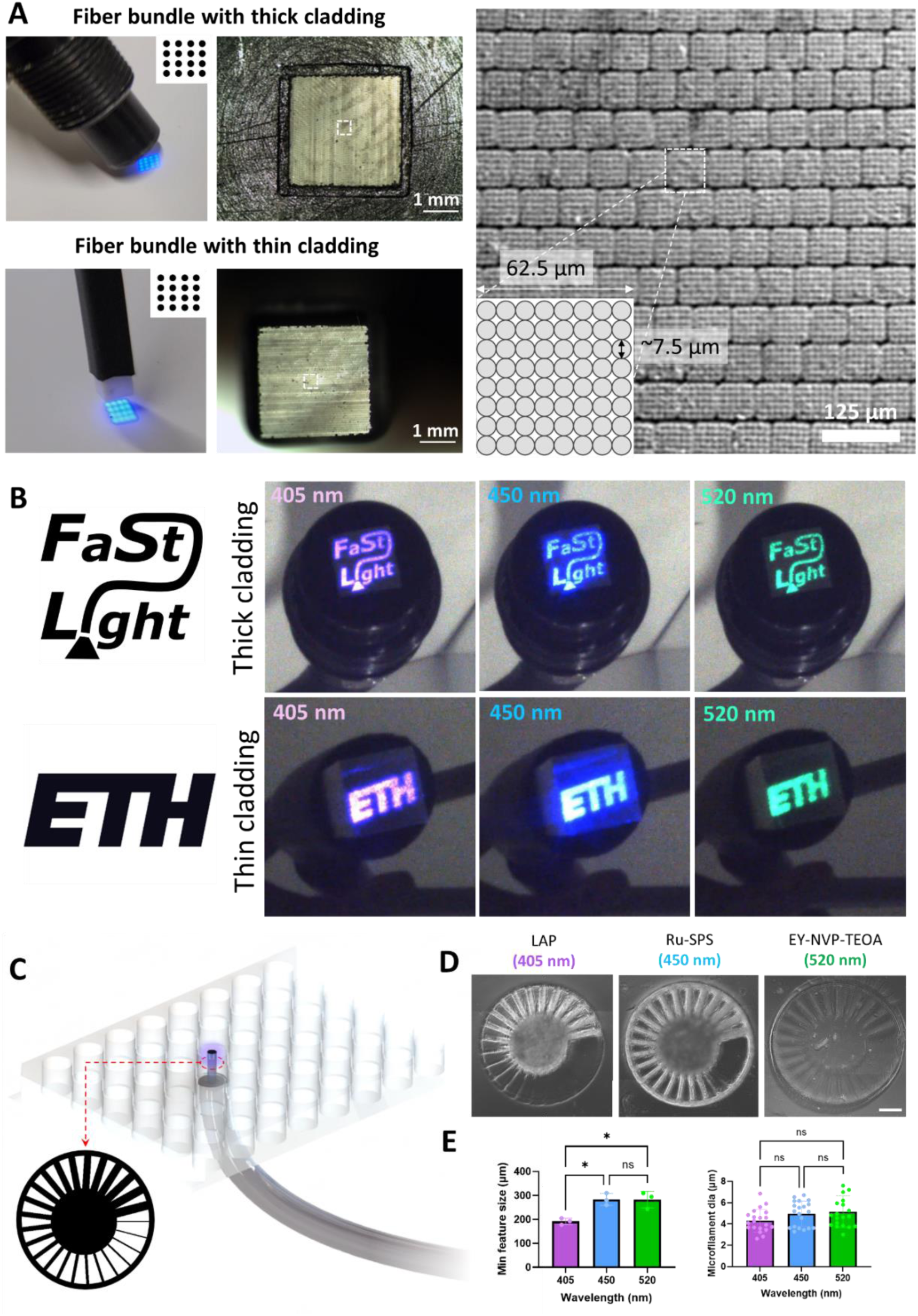
Different types of image guide fiber bundles in the FaSt-Light apparatus. **A.** Fiber bundles with thick and thin cladding. Inset images (dotted rectangles) show the corresponding building blocks of the fiber arrays. **B.** Different images (FaSt-Light and ETH logos) projected at different wavelengths. **C.** Fiber bundles being used for bottom-up projection of spoke wheel image (Φ = 3 mm) into the resin contained within 48 well plates. **D.** Crosslinked spoke wheel constructs made at each wavelengths (scale bar: 500 µm). **E.** Minimum attainable feature size (i.e., spoke width) and microfilament diameters at each wavelength. * represents p < 0.05.

To fabricate the constructs for resolution measurements, we filled GelMA resin (100 µl per well; corresponding to ∼1.34 mm height) in 48 well plate and projected spoke wheel images (Φ = 3 mm) from fiber bundle positioned underneath the well plates (illustrated in **Figure 3C**). Of note, light intensity attenuates as it travels through the fiber bundle. For instance, the light intensity at the output of the projection apparatus (i.e., the input of the fiber bundle) at 405, 450, 520 nm was 31.1, 34.5 and 40 mW/cm^2^, respectively. While the power at the output of the fiber bundle (3×3 mm^2^ section and 75 cm long; with thick cladding) for 405, 450, 520 nm was 14.1, 16.7 and 22.5 mW/cm^2^, respectively. Accordingly, we increased the exposure duration of the different GelMA formulations at each wavelength to achieve the same light dose (in mJ/cm^2^; Figure S1) for material crosslinking. The spoke wheel prints at each wavelength within the 48 well plates are shown in **Figure 3D** (actual setup is shown in Figure S2; also see Video S3 for the spoke wheel projections at each wavelength from the fiber bundle), and the corresponding analysis of the minimum attainable feature size and the microfilament diameters are shown in **Figure 3E**. While the microfilament diameters in the three different photoresin formulations crosslinked by the fiber bundle (images of microfilaments are shown in Figure S3) were not different, the minimum attainable feature size for the resins containing LAP (i.e., 405 nm) was the lowest (i.e., highest resolution), while the resins containing Ru-SPS (i.e., 450 nm) and the EY-NVP-TEOA groups demonstrated the higher feature sizes (i.e., lower resolution). The reduced resolution in resins containing Ru-SPS can be attributable to the increased tyrosine crosslinking (similar to **Figure 2D**), the reduced resolution in resins containing EY-NVP-TEOA can be attributed to the prolonged exposure times (71 s) required by the resin formulations. In contrast, the resins containing LAP or Ru-SPS only needed 26 or 10 s for the fiber-assisted crosslinking of the resin, respectively. A prolonged exposure can also lead to more non-specific crosslinking due to the diffusion of free radicals within the photoresin, thereby affecting the resolution. Despite the prolonged exposure needed for the formulations containing EY-NVP- TEOA, it had the lowest elastic modulus (∼ 0.9 kPa; see **Figure S4**), whereas the moduli for formulations containing Ru-SPS and LAP were comparable (∼ 2 kPa). For all subsequent tests, we only used the resin formulations containing LAP as they resulted in the highest resolution amongst the three formulations (Figure 3E). Furthermore, despite the faster print times enabled by the resin formulations containing Ru-SPS, the formulations containing LAP are simpler (i.e., a single photoinitiator component) and have lower attenuation coefficient,^[21]^ resulting in greater penetration of the light into the photoresin. To characterize the depth of crosslinking, we only compared the resin formulations containing LAP or Ru-SPS (**Figure S5**), as these formulations resulted in similar elastic moduli. In these tests, we projected custom images using the image guide fibers into 10 mm path length cuvettes. Here, while photocrosslinking was achieved up to 9 mm depth in the formulations containing LAP, those containing Ru-SPS only demonstrated photocrosslinking only up to 2.5 mm in depth (Figure S5). In these tests, the optical self-focusing effect, where the light is preferentially guided into the crosslinking polymer due to changes in refractive index,^[16,21]^ contributed to a reduced scattering of light and maintenance of image features through the crosslinked constructs.

In addition to different types of cladding, the FaSt-Light apparatus can also deploy different sizes of fiber bundles to accommodate the projection of larger images into the photoresins. For instance, we compared two different sizes of fiber bundles – 3×3 mm^2^ and 4×4 mm^2^ – coupled to the projection apparatus (**Figure 4A**). While the overall sizes of the bundles were different, the constitutive blocks of fiber arrays in the two bundles were the same (representative image in **Figure 3A**), but the number of such blocks is larger in the larger fiber bundle. To compare the effects of the different sizes of fiber bundles, we projected the same spoke wheel images (ɸ = 3 mm) using the two fibers within 48 well plates as previously shown in Figure 3C. As discussed previously, only photoresin containing LAP was used in these experiments. The representative print and the microfilament distribution within the crosslinked photoresin using the two different fiber bundles are shown in **Figure 4B**. The corresponding analysis of the print resolution and the microfilament distribution (**Figure 4C**) revealed no difference between the two fiber sizes. This is expected, since the constitutive blocks of the fiber arrays within the two bundles are of the same size. Therefore, the pixel size of the projected images (same as the size of individual fibers; ɸ = 7.5 µm) are the same through the two fibers.

**Figure 4.**
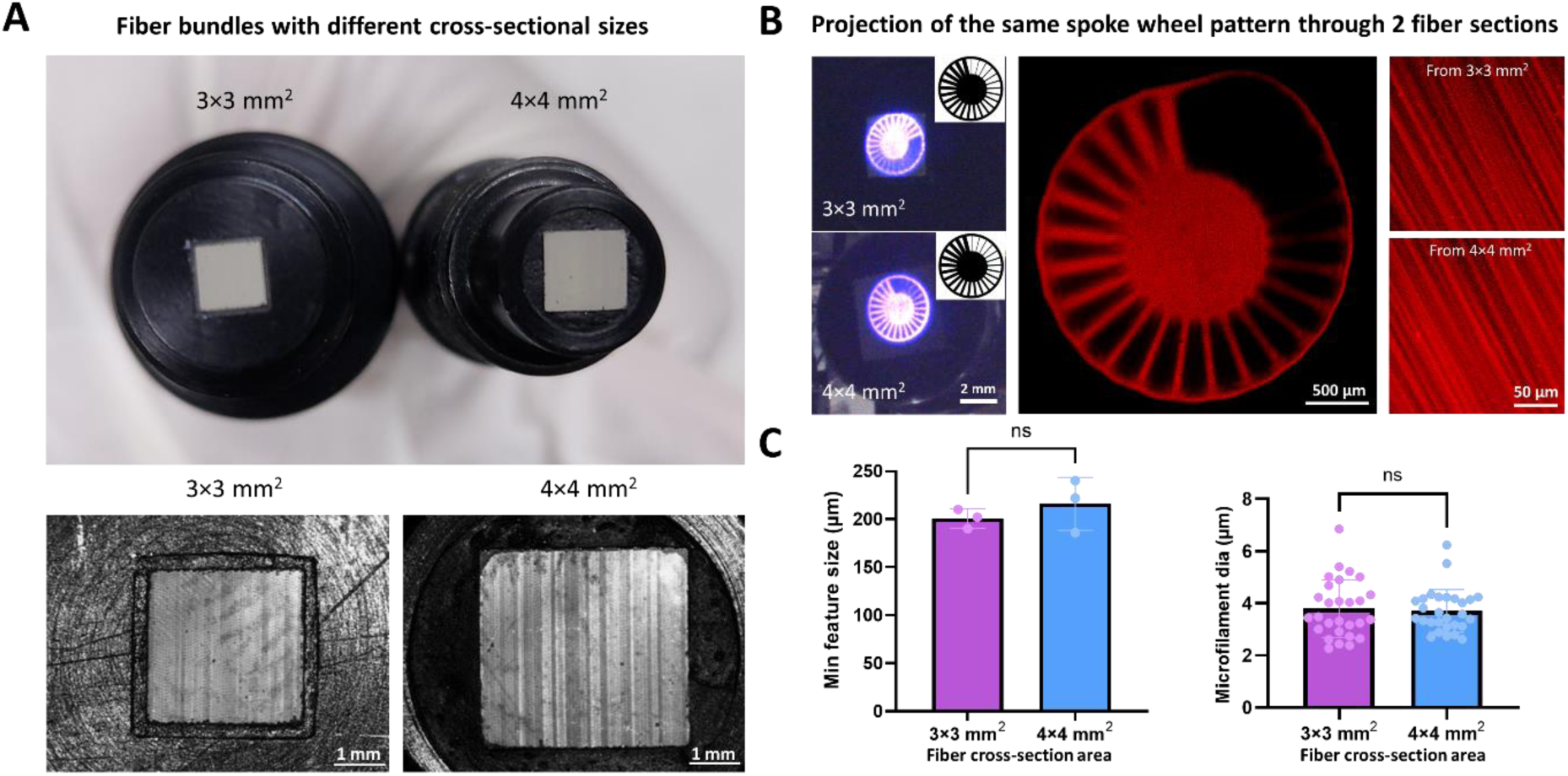
Comparison of different sizes of fiber bundles in the FaSt-Light apparatus. **A.** Two different fiber sections (3×3 mm^2^ and 4×4 mm^2^) were compared, each of which constituted the same blocks of fiber arrays (each block is an 8×8 grid of individual fibers), but the number of blocks were larger in the 4×4 mm^2^ fiber bundle. **B.** Projected spoke wheel image (ɸ = 3 mm) from the two different fiber bundles and the crosslinked photoresin and its microfilament distribution. **C.** The corresponding results of minimum feature size and microfilament diameter.

In applications involving imaging in hard-to-reach regions (e.g., endoscopy), the fiber bundles are typically located a few mm or cm away from the target object to allow effective imaging and avoid contact with the object.^[35,36]^ In these applications, custom lens arrangements are often coupled to the fiber bundles for capturing a wider area.^[35,36]^ Accordingly, for in situ printing applications, we posit that the fibers will be located a few mm to cm away from the photoresins, and custom lenses could be used for projecting wider images onto the resins. Therefore, we observed the features of the projected images at different distances from the fiber bundles to characterize any loss in resolution. We also compared a bi-concave lens (dioptre = -111.1 m^-1^; half-angle divergence = 16.7°) coupled at the output of the fiber bundle to be able to widen the projected image. For these experiments, a white screen (0.1 mm thick) was moved away from the fiber bundle and a charge coupled device (CCD) camera was positioned at the opposite end to capture the images being projected into the screen (illustrated in **Figure 5A**). The images captured at different distances from the fiber without the bi-concave lens have been shown in **Figure 5B**, and those with the coupled bi-concave lens (setup illustrated in **Figure 5C**) have been shown in **Figure 5D**. While the projection at the output of the fiber remains collimated up to 3 mm away from the fiber (i.e., the image size and features remain the same), the image gradually expands and loses its features as the distance increases (**Figure 5E**). With the bi- concave lens allows expansion of the projected image. However, the image features are only maintained up to 3 mm away from the fiber (**Figure 5E**). In both conditions (i.e., with or without bi-concave lens), the intensity of the image normalized to the surface area also gradually decreases (**Figure 5F**). We further plotted the variations in the gray value of the intensities across a section of the image at different distances (**Figure 5G**). Here, for both the fiber bundle without or with bi-concave lens, the distance between the consecutive maxima (and minima) of the gray values increases after 3 mm, showing the increased distribution of light intensities as the distance increases more than 3 mm. Plotting the gray value distribution allowed us to quantitatively assess the shape index of the images, where the distance between the maxima was divided with the feature size of the image had it been a lossless projection (additional details in methods). The shape index results (**Figure 5H**) also support the qualitative assessments, where the index is above 0.9 (i.e., a high fidelity projection) for up to 3 mm distance from the fiber, but decreases as the distance progresses. As a potential application, we show that the images can be projected from a distance of 3 mm onto resin (GelMA with LAP) filled into skin defects (∼ 1 mm deep) in a rat cadaver. The FaSt-Light projection and the corresponding high-fidelity prints of the window pattern onto rat skin defects are shown in **Figure 5I** (also see Videos S4 and S5). Here, when fiber with or without lens was used, the duration of the projection was fine-tuned to be able to achieve a cumulative light dose of 370 mJ/cm^2^ (optimal for LAP; Figure S1).

**Figure 5.**
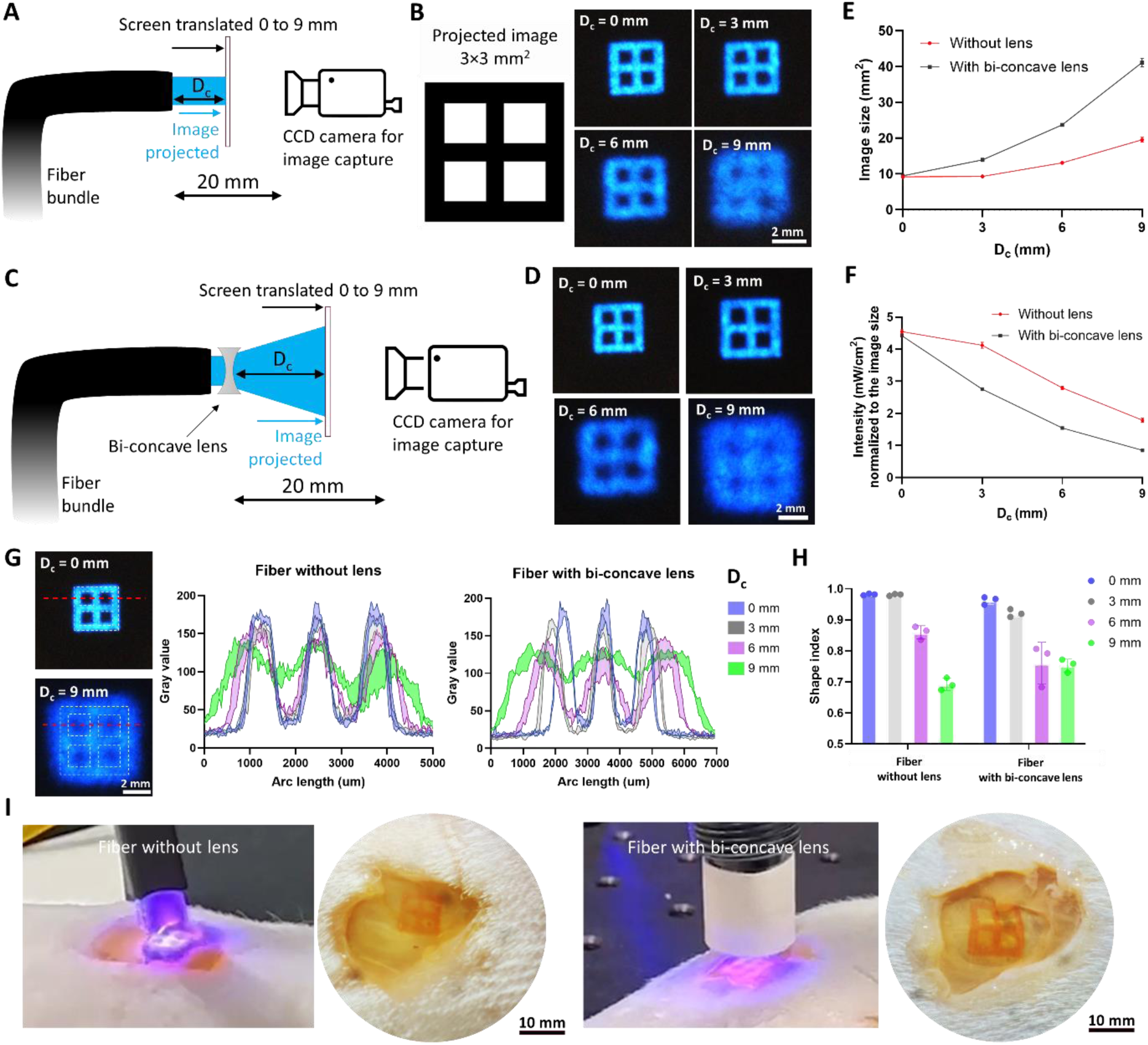
Characterizing the loss in resolution in the images projected from the fibers with or without bi-concave lens. **A.** Setup used to capture the images projected from the fiber bundle onto a screen at different image capturing distances (D_c_ = 0 to 9 mm). The distance between the output of the fiber bundle and the camera is fixed to 20 mm. **B.** The projected images of a 3×3 mm^2^ window pattern through the fiber. **C.** Setup to capture images projected at different distances from a fiber coupled with a bi-concave lens. **D.** Projected images from the lens-coupled fiber. **E.** Size of the image projected from the fiber. **F.** Intensity normalized to the size of the projected image from the fiber. **G.** Estimation of the gray value along the red dotted line in the projected image. White dotted line represents the outline of the ideal projection image. **H.** Shape index of the projected image when compared to the intended image (after accounting for the expansion from the lens), at different distances from the fiber without or with the bi-concave lens. **I.** Demonstration of the use of fibers for projecting images onto resin filled in a skin defect on rat cadavers. Here, the fiber without or with bi-concave lens was held at a distance of 3 mm from the resin-filled defect and the window-shaped pattern was projected onto the resin. The camera images show the crosslinked resin after washing.

While the above tests demonstrated that the fiber bundles could be used either standalone or coupled with a bi-concave lens to project images onto the resin, the distance of the fiber from the resin was limited to a maximum of 3 mm to achieve a high-fidelity projection. However, for applications where larger area needs to be covered (e.g., tissue defects spanning several cm), a small distance limits the area of the resin which can be effectively crosslinked using a single projection. Here, we demonstrate that a larger projection and resin crosslinking could indeed be achieved by coupling with fiber bundle with a C-mount lens (**Figure 6A**), which allows one to preserve the fidelity of the projected image. C-mount lenses consist of a custom assembly of optical components which enable high-resolution imaging in scientific and industrial applications.^[42,43]^ **Figure 6B** shows high fidelity image projection, from the C-mount lens coupled fiber bundle, for all three wavelengths at a distance of 20 mm from the screen. To compare the projection image size and fidelity with the previous results in **Figure 5B**, we projected the window-shaped image (3×3 mm^2^) and translated the screen from 0-9 mm (**Figure 6C**). The corresponding projection images at the different distances from the C-mount lens are shown in **Figure 6D**. Similar to the bi-concave lens, the C-mount lens allows expansion of the images, and hence a reduction in the normalized light intensity (**Figure 6E**). However, unlike the bi-concave lens, the shape fidelity of the projection image is maintained, which is evident from the results of the gray value profiles and corresponding analysis of the shape index when comparing the actual image to the intended image (**Figure 6F**, details in methods). Here, the shape index is >0.9 for distances greater than 0 mm, which is highly desirable as most applications will require the fiber bundles to be located away from the target resin. To demonstrate the use of C-mount coupled fiber bundle we created a 15 mm wide skin defect (∼1 mm depth) in a rat cadaver and positioned the C-mount lens 9 mm above the skin defect. Referencing the image expansion from **Figure 6D**, a 3×3 mm^2^ projected image expanded to ∼9×9 mm^2^ (i.e., a 3X magnification) at a projection image at a distance of 9 mm. Accordingly, we projected a 5×5 mm^2^ meshed image from the fiber bundle to be able to cover the entire skin defect (**Figure 6G**). After filling the defect with the photoresin (GelMA with LAP), we projected the meshed image pattern from the fiber bundle. Here, as the intensity lowered due to the expansion of the projected image, the exposure duration was adjusted to achieve the same light dose required from crosslinking (similar to the experiments shown previously in Figure 5I). The corresponding crosslinked mesh after washing the defect region with PBS is shown in **Figure 6G** (lower panel). In the discussion section, we have highlighted the potential regenerative applications of the current state of the C-mount-coupled FaSt-Light apparatus, and further discussed potential approaches to reduce the form-factor of the lens-coupled fibers to facilitate minimally invasive image delivery for crosslinking resins over hard-to-reach tissues in an in vivo application.

**Figure 6.**
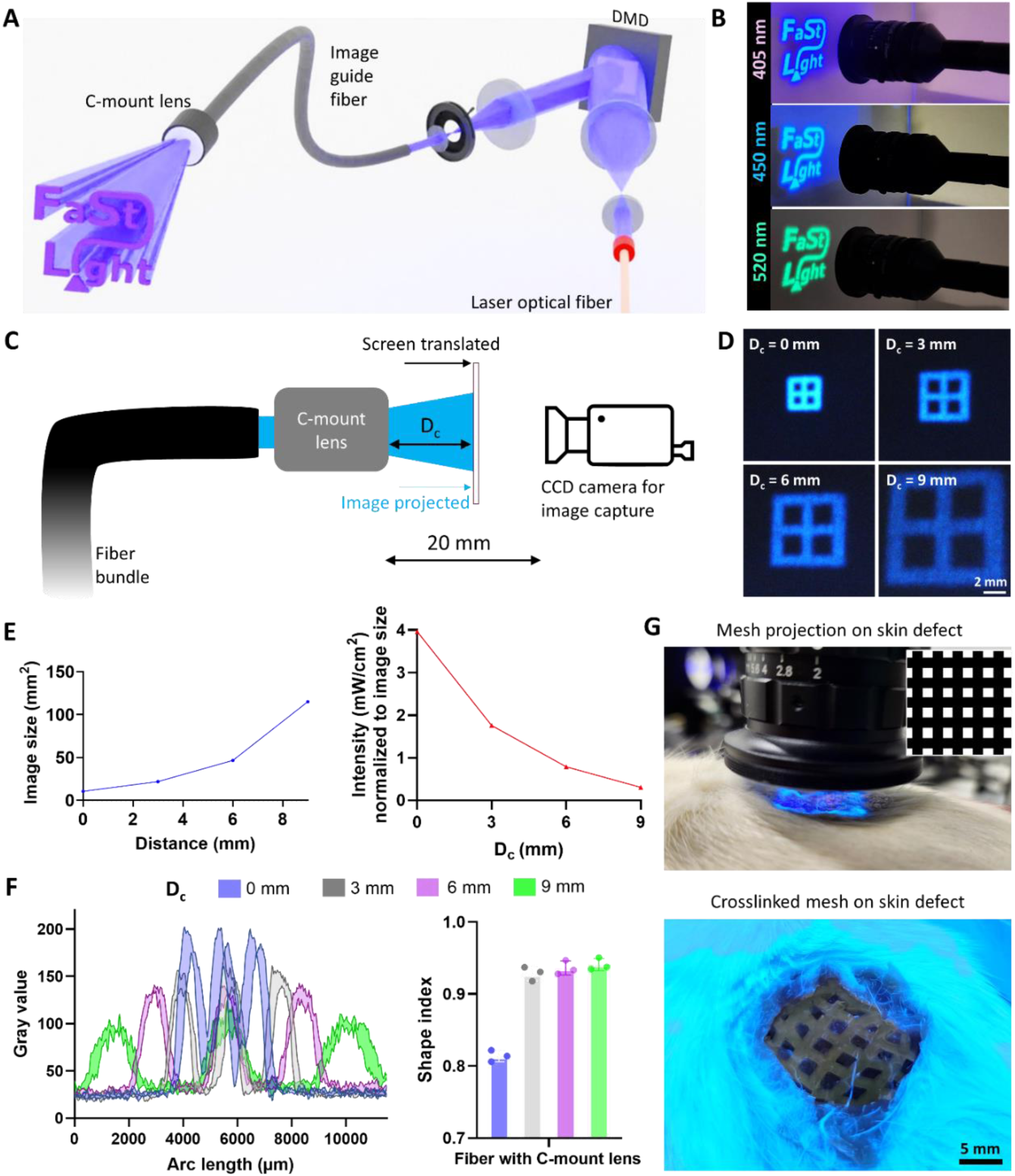
Demonstrattion of high fidelity image projection using fiber bundles. **A.** Concept image of coupling of C-mount lenses onto fiber bundles to preserve image fidelity. **B.** Projected FaSt-Light logo at each wavelength at a distance of 20 mm away from a white screen. **C.** Setup for capturing the images projected onto a screen from a fiber coupled with C-mount lens. **D.** Images projected onto a screen at varying distances (0-9 mm) from the lens. **E.** Variation of the image size and the light intensity normalized to the image size at varying distances from the lens. **F**. Variation of the gray values at increasing distances from the C-mount lens and the corresponding shape index when comparing the projected images to the intended image after accounting for the expansion from the lens. **G**. Crosslinking of resin (inset in the top panel shows the 5×5 mm^2^ mesh image which was used) over a rat skin defect. The bottom panel shows the crosslinked mesh (GelMA with LAP was crosslinked at 405 nm and later washed using PBS) over the skin defect.

Given that the fiber bundles allow maintenance of the light structure (macroscopic image and microscopic speckles), the FaSt-Light apparatus can be an effective tool for the in situ biofabrication for tissue repair (e.g., defects of muscle, tendon, nerves, etc.). To demonstrate the use case for muscle biofabrication, we encapsulated C2C12 myoblasts at 10^6^ cells/ml within the GelMA formulations with LAP and transferred the resins within 10 mm path length cuvettes. Of note, in these studies, we had supplemented the resin with iodixanol (a biocompatible refractive index matching agent) to reduce the light scattering effect due to the presence of cells (additional details in methods).^[12,20]^ Also, the light dose was increased by 20% to mitigate the effects of light attenuation due to presence of cells. After thermo-reversible gelation at 4°C, we used the fiber bundles to project rectangular images (1×3 mm^2^) into the cuvettes (**Figure 7A**). Due to the optical self-focusing effect, the light projection could crosslink the resin along the entirety of the cuvette, creating 10 mm long rectangular sheets where the microfilaments were present across the entire length (**Figure 7B**). Since the LAP photoinitiator absorbs light, thereby affecting the light dose along the length of the cuvette, we further characterized the differences in maturation along three regions – region near the fiber bundle during biofabrication, middle zone and away from the fiber bundle. After maturation (1 week of growth + 1 week of differentiation), the muscle constructs (**Figure 7C**) in each of the three regions exhibited multinucleated myotubes which stained positively for sarcomeric alpha actinin. Notably, we did not observe changes in the three regions with regards to the myotube diameter (**Figure 7D**) and density (**Figure S6**), which is likely due to the short duration of culture to be able to observe region-specific changes. While the muscle constructs also exhibited sarcomere structures resembling native mouse muscle (∼2.2 µm; **Figure 7E**),^[44]^ the myotube diameters are smaller than those found in native mouse muscle tissues (∼50 µm),^[45]^ which is likely due to the short maturation period, low cell densities and absence of other cell types (e.g., fibroblasts and endothelial cells, etc.) supporting muscle maturation.^[46,47]^ Longer culture, higher cell densities and co-culture systems will be investigated in our future work to achieve biomimetic myotube diameters and other attributes (synchronous contractility, connective tissue deposition, neovascularization, etc.) resembling native muscle tissue.^[46,47]^

**Figure 7.**
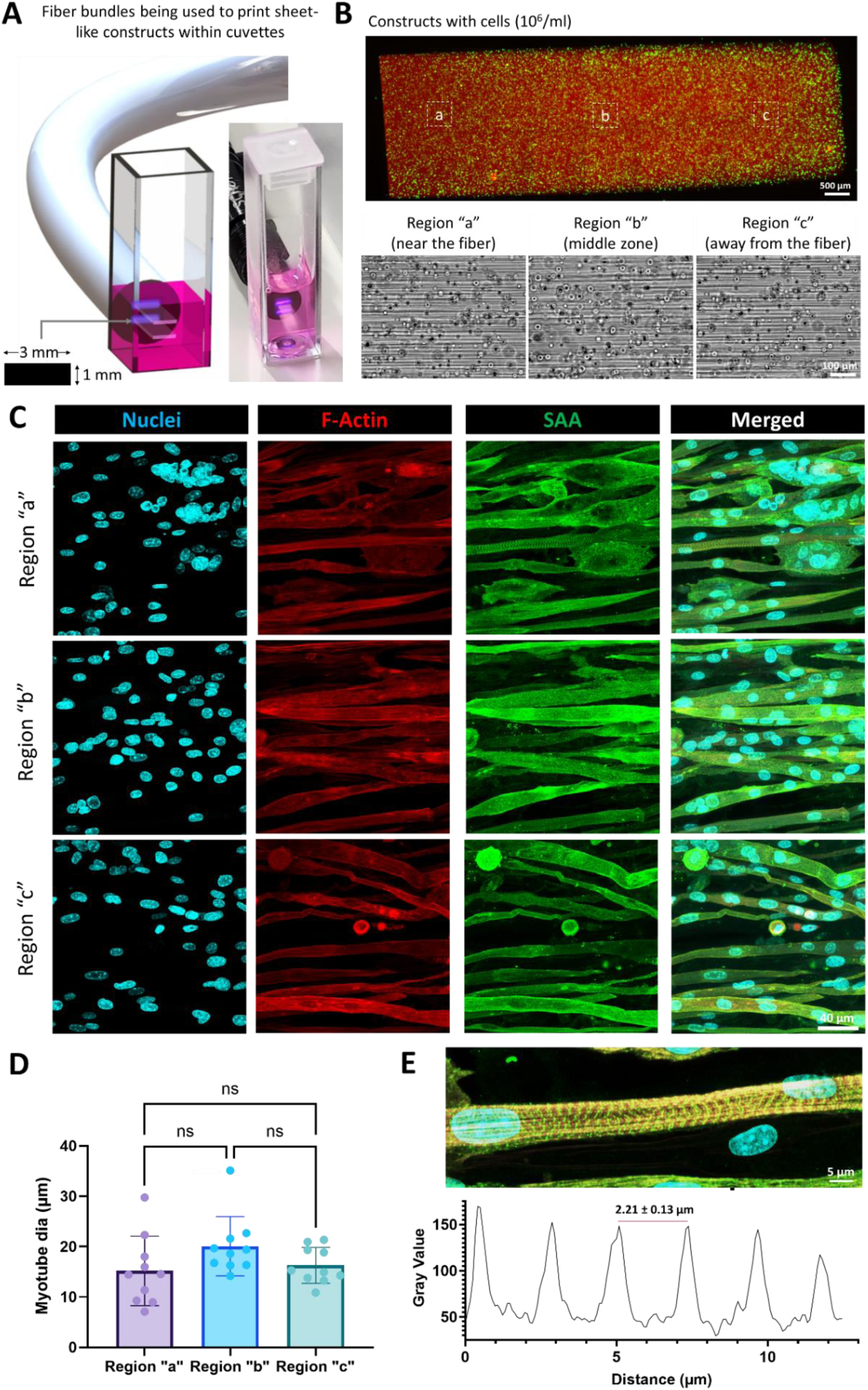
Use of Fast-Light projection for fabrication of muscle constructs. **A.** The muscle constructs were fabricated by projecting rectangular images (1×3 mm^2^ at 405 nm) from image guide fibers in contact with 10 mm path length cuvettes containing the photoresin (GelMA + LAP). The photograph used rhodamine-labeled GelMA to allow better visualization. **B.** 10 mm long sheet constructs with encapsulated C2C12 cells (labeled green using Calcein AM dye; red portion is the rhodamine-labeled GelMA). The images for the insets (“a”, “b” and “c” regions) are shown in the bottom, which demonstrate the presence of microfilaments around the cells in each region. **C.** Matured constructs (1 week of growth + 1 week of differentiation) exhibit key muscle sarcomeric alpha actin staining. Scale bar is 40 µm. **D.** The myotube diameter is not different in the three regions of the constructs. **E.** Regularly interspersed sarcomere structures are observed in the constructs with an average sarcomere spacing of 2.21 µm.

In addition to in situ biofabrication of cellularized constructs, the fiber bundles can also be used for fabricating acellular biomaterial grafts in situ for new tissue formation and repair. Here, we show that the presence of microfilaments can provide topographic cues for cell guidance and infiltration. As a demonstration, we encapsulated myoblasts (C2C12; 10^6^ cells/ml) in GelMA resins (with LAP; labelled with rhodamine for clear distinction between the layers) and filled the resins within 8 well plates (**Figure 8A**) to create a layer height of ∼1 mm (100 µl of resin was added). The resins were then crosslinked using bulk light illumination (405 nm) in a UV box (see methods for details), and a scalpel was used to remove half of the crosslinked cell-laden hydrogel. The constructs were then thoroughly washed with PBS (to remove any uninitiated LAP), and the excess PBS was removed. In the remaining portion, fresh acellular GelMA was added and crosslinked using the fiber bundle (plain image was projected from the fiber) from the side of the well plate (**Figure 8B**). The resulting crosslinked constructs (**Figure 8B**) feature two regions – a rhodamine labelled region with encapsulated cells (bulk light crosslinked), and an acellular region featuring microfilaments (crosslinked using the fiber bundle). Of note, we also observed some microfilaments in the cellular region which was crosslinked with bulk light. This is likely due to diffusion of LAP from the freshly added resin which was then crosslinked with the fiber bundle which introduces the microfilaments. After supplementing media over the crosslinked constructs in the well plates and culturing over 96 h (media changes every 24 h), we observed cell infiltration into the constructs (**Figure 8C**) up to 3.08 ± 0.41 mm and a predominantly aligned cell morphology throughout the constructs, which demonstrates suitability of the microfilamented constructs in effectively guiding cell infiltration. We further verified whether such an approach could be deployed within an in vivo wound muscle defect in rat cadavers (concept illustrated in **Figure 8D**). For these experiments, we created an incision in the back of rat cadavers and created a 5×3×3 mm^3^ defect in the dorsal muscle tissue. Here, a larger incision allowed better visibility of the use of fiber bundle for resin crosslinking. The fiber bundle and a syringe needle were inserted from two separate incisions on either side of the wound site, and the resin dispensed from the syringe followed by blank image projection from the fiber bundle (**Figure 8E**). Here, the excised tissue allowed a better view of the crosslinked gel in the defect (Figure 8E). When the crosslinked gel was extracted from the tissue site using a spatula and imaged, the microfilaments were prevalent throughout the entire length of the defect. Considering the results from in vitro tests showing cell infiltration, the in vivo proof-of-concept experiments demonstrating the fabrication of microfilamented hydrogel scaffolds in can open new applications for the regeneration of musculoskeletal defects. In the discussion, we further highlight the future developmental work on the FaSt-Light approach and the regenerative applications this approach can be used for.

**Figure 8.**
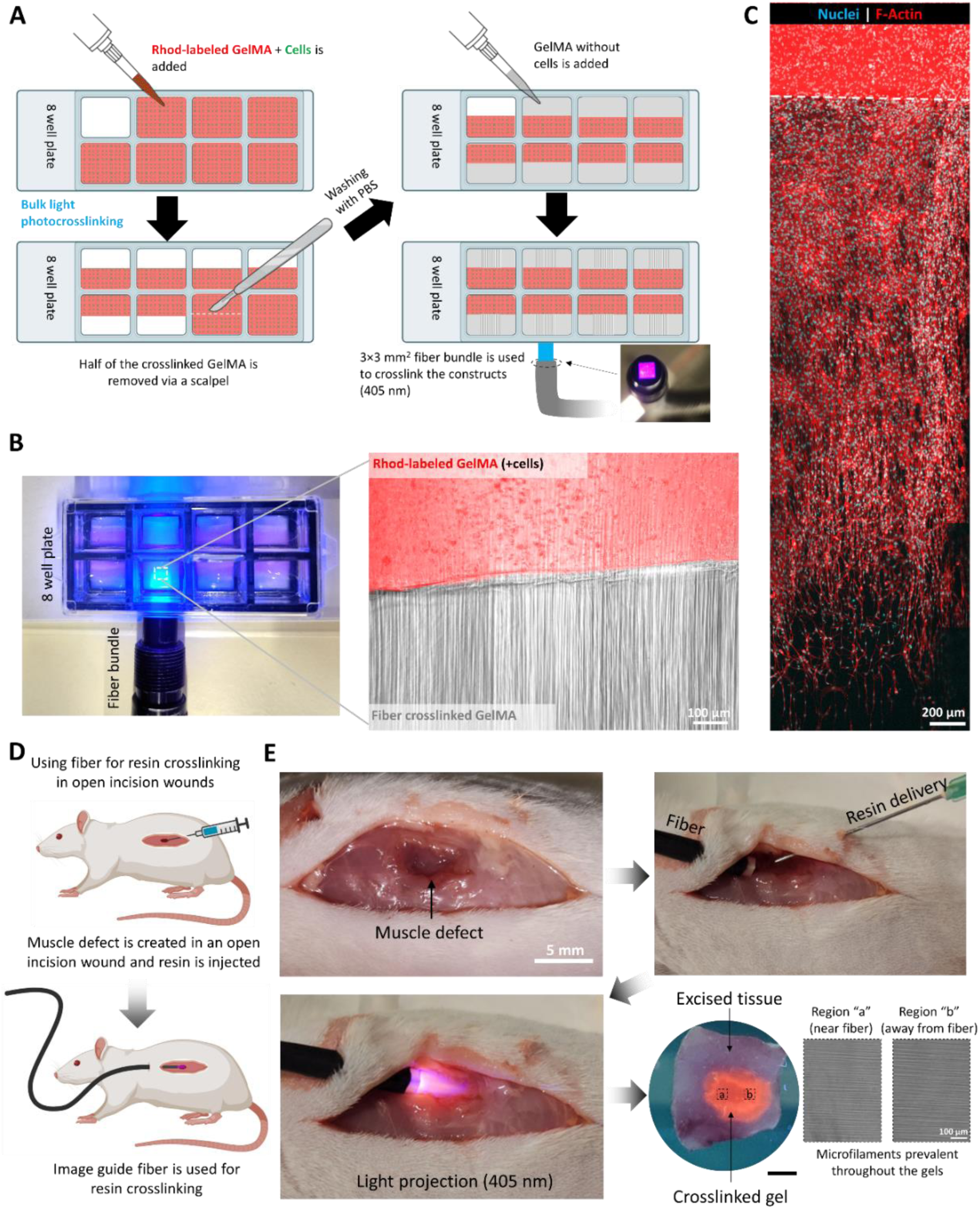
Concept experiments demonstrating cell infiltration into acellular microfilamented matrices, and the use of FaSt-Light approach for in vivo resin crosslinking. **A.** Fabrication of the constructs featuring Rhodamine-labeled GelMA (crosslinked with bulk light exposure to 405 nm in a UV box) containing cells, and acellular GelMA crosslinked with the image guide fiber. **B.** Camera image (left) of the image guide fiber to crosslink the acellular GelMA to the cellular GelMA and the micrographs showin cellular (rhodamine-labeled) and acellular (unlabeled) regions of the constructs. **C.** Stitched micrographs showing cell infiltration into the acellular microfilamented GelMA region 3 days after culture (dotted lines represent the border between the different GelMA regions). **D.** Concept experiments on the use of FaSt-Light apparatus to achieve microfilamented resin crosslinking within a muscle defect in rat cadavers. **E.** Camera images of the muscle defect (incision kept larger than the defect to allow easy visualization), and the insertion of the fiber bundle and syringe needle via separate incisions on either side of the defect. After the light projection, the tissue is excised and imaged. The crosslinked construct is further extracted from the tissue and microstructure (after imaging) shows the prevalence of microfilaments throughout the crosslinked gel.

## Discussion

Image guide fiber bundles have traditionally been used for capturing images from hard-to-reach regions such as during endoscopy, and to guide the images to camera sensors. We have demonstrated a reverse approach called FaSt-Light, where the bundles were used to guide the images generated from a projection system. The approach offers projection of images at multiple wavelengths, allowing the crosslinking of resins along custom shapes, which is not permissible by other approaches which rely on light guide fibers projecting homogeneous light beams. Here, we highlight the current state of development and future scope which could allow the FaSt-Light approach to develop into a highly scalable and flexible apparatus for resin crosslinking, potentially transforming the field of minimally-invasive in situ biofabrication.

For the fiber bundles without or with bi-concave lenses used this work, the projected images lost their fidelity for distances greater than 3 mm. Here, C-mount lenses allowed maintenance of projected image fidelity for larger distances, and that the widening image lost intensity which needed compensation in terms of prolonged exposure duration to achieve resin crosslinking. However, the form-factor of the C-mount lenses was large, rendering such a system less suitable for minimally invasive photo-bioprinting applications. Nevertheless, in its present state current setup could be used for resin crosslinking onto open tissue sites such as diabetic chronic wounds,^[48,49]^ or over the myocardium (i.e., applications involving direct in situ cardiac patch fabrication) during open heart surgery,^[50,51]^ etc. Here, the aspects of microfilaments guiding native cell infiltration, differentiation and new tissue formation would be interesting to study and are a potential exploration area in the future. In such applications, the resin viscosity will have to be carefully fine-tuned for ease of application and conformation of the resins to the complex contours and dynamic movements of the underlying tissue (i.e., the resin should not flow away due to the tissue shape or its movements). Since the GelMA resins in the present work undergo thermo-reversible gelation under colder temperatures (typically < 24°C), the present resin formulations can also be potentially suited to in situ biofabrication in dynamic environments, where one could directly add the resin at a cooler temperature and then quickly photocrosslink it before it liquefies.

Notably, for applications involving side-ways projection of images into the resins, we observed that the optical self-focusing effect enables the crosslinking of constructs up to 10 mm in length, which indicates that the fibers as standalone devices could potentially be used for the *in situ* fabrication of aligned tissue constructs or grafts which can promote cell infiltration. Potential applications include the treatment of volumetric muscle loss or peripheral injury repair, etc.^[52–55]^ However, further investigation is needed as to the extent of the optical self-focusing effect based on the photoinitiation system. Since LAP has a lower attenuation coefficient compared to other initiation systems such as ruthenium complexes,^[21]^ it can be better suited for applications requiring larger defects. However, whether LAP-based resins can allow resins crosslinking for defects spanning several centimetres will be a scope of future investigation. Furthermore, LAP is used primarily with modified resins such as those relying on acrylate, methacrylate, or thiol-ene chemistries, etc.^[56,57]^ This, in conjunction with under-studied immunogenic effects of LAP, affect the translational potential of the resins. In contrast, photoinitiators such as ruthenium complexes (e.g., Ru-SPS system) or riboflavin phosphate (vitamin B-12) can allow crosslinking of natural resins such as collagen or fibrinogen using the native tyrosine or histidine residues.^[58,59]^ Such a modification-free crosslinking approach simplifies the formulations, improves the cost effectiveness and the translational potential of these resins. The low penetration of light in these photoinitiation systems can still be relevant for applications involving the fabrication of patches (e.g., for wound healing or cardiac patches).^[8]^ Even for large scale defect-filling, to compensate for the limited penetration depth of crosslinking, the fiber bundles could be made a part of toolheads which can dispense the photoresins over the tissue site concomitantly with the image projection from the fibers. Here, the fiber can be made to mechanistically transverse (e.g., through a robotically guided toolhead) from one side of the defect to the other for consistent crosslinking throughout the entire defect. Here, photoabsorbers would be necessary to prevent overcrosslinking in the already-crosslinked regions.^[21]^ Alternatively, resin crosslinking in larger defects can also be executed using two fibers from either side of the defect. Here, accounting for the attenuation of light in the resin, the light doses will need to be carefully calibrated to achieve a cumulative light dose at the centre of the construct equalling that on the extremities of the constructs, to be able to achieve constituent mechanical properties throughout the crosslinked resin. Importantly, washing off the uncrosslinked photoresin may also be needed to mitigate any unwanted immunogenic effects of the remnant photoinitiators or unreacted monomers.

The aspect of being able to project multiple wavelengths allowed us to demonstrate the use of different photoinitiation systems with the FaSt-Light apparatus. However, the projections were executed one wavelength at-a-time. Notably, the DMD chip used in the present work has high frame refresh rates (9523 Hz), which can allow synchronization of the image being projected to the laser wavelength being activated. This can allow the projection of spatial regions featuring different wavelengths within a single projection image. Such a system, when effectively combined with photocrosslinking or photodegradation chemistries which rely on different wavelengths, can allow in situ photo-bioprinting of multi-material constructs.^[2,60]^ Here, the photoinitiator will have to be carefully selected, as initiators such as the Ru-SPS have light absorption at wide range of wavelengths (see Figure S7 demonstrating the Ru-SPS initiated tyrosine crosslinking of pristine collagen formulations at each wavelength).

In the present work, we have demonstrated that the FaSt-Light approach can be used for the fabrication of anisotropic muscle tissue constructs or acellular grafts which promote cell infiltration and alignment. We have also demonstrated proof-of-concept studies on the in vivo fabrication of patch-like geometries or microfilamented biomaterial grafts in rat cadavers. Verification and validation of the suitability of FaSt-Light approach towards applications such as wound healing,^[61,62]^ volumetric muscle loss,^[52,53]^ peripheral nerve injury repair,^[54,55]^ etc., would require detailed in vitro characterization studies with multiple cell types and in vivo studies in live animal models. For these injury models, careful consideration will be needed in terms of the orientation of microfilaments (i.e., the direction of light projection from the fiber bundles), which will ideally need to match that of the tissue at the distal and proximal ends of defect to enable effective cell infiltration and matrix deposition which aligns with the native fiber orientation. Also, the stiffness of the in situ photo-fabricated grafts will need to be carefully tuned as it plays a crucial role in cell inflammatory responses, tissue healing and scar formation.^[63]^ In the future, we envision the integration of the FaSt-Light system onto mechanised robotic toolheads which can maneuver into hard-to-reach regions and precisely position and perform in situ printing of multicellular and multimaterial grafts at the tissue site. Mechanisation of the fiber bundle-assisted printing approach with new advances in deep learning of the images captured during the biofabrication process (i.e., image guided therapy) can further improve the translational potential for an in vivo usage by reducing human errors.^[64,65]^

## Conclusion

FaSt-Light integrates image guide fiber bundles with a multi-wavelength projection system, offering a new approach for in situ photo-biofabrication. The images are guided through fiber bundles which come in a variety of shapes and sizes, offering flexibility on the maneuverability, handleability and the location of fabrication. The use of multiple wavelength projections allows compatibility with a variety of photoinitiation mechanisms. In addition to a controlled microarchitecture (through controlling the projected image), the resins crosslinked with FaSt-Light also feature microfilaments (due to optical modulation instability) which allow cell-guidance and infiltration, opening new applications for in vivo tissue biofabrication or repair. By the coupling of appropriate lenses to the fiber bundles, the image fidelity can be maintained for image projections several centimetres away. The FaSt-Light process offers new pathways of research into tools which combine multiple printing modalities, mechanisation and concomitant imaging with printing, potentially ushering in a new era of in situ biofabrication and transitioning away from traditional benchtop devices.

## Methods

### Hardware and consumables sourcing

The fiber bundles were procured from Schott AG (Mainz, DE). The lenses and optomechanical components for the FaSt-Light apparatus, as well as the power meter (S121C; 400-1100 nm measuring range) were procured from Thorlabs GmbH (Bergkirchen, DE). The DMD (evaluation board DLPLCR67EVM) and its controller (DLPLCRC900DEVM) were procured from Texas Instruments. The multi-wavelength laser light engine was procured from Lasertack GmbH (Fuldabrueck, DE). 10 mm path length cuvettes were procured from Thorlabs GmbH (CV10Q35F). 8 well plates for cell infiltration tests were procured from ThermoFisher (Nunc™ Lab-Tek™ II Chamber Slide™ System; 154534). UV box for fabricating bulk GelMA resins (BSL-01) was procured from Opsytec Dr. Gröbel GmbH.

### Chemical sourcing

Merck KGaA, (Darmstadt, DE) was the source to procure Gelatin (Type B, G6650) and methacrylic anhydride (760-93-0), acryloxyethyl thiocarbamoyl rhodamine B (Rhod-Acr; 908665), Dulbecco’s modified eagle medium (DMEM; D6429), Penicillin-Streptomycin (P/S; P4333), Trypsin-EDTA solution (T4049), fetal bovine serum (FBS; F1283), horse serum (H0146), propidium iodide (537059), Calcein-AM (148504-34-1), Triton-X100 (T8787), paraformaldehyde (PFA; 148504-34-1), bovine serum albumin (A7030), and phalloidin (P1951). Hoechst (33342) used for nuclear staining was procured from ThermoFisher. The primary antibodies consisted of Monoclonal Anti-α-Actinin (Sarcomeric) antibody (SAA) produced in mouse (A7811, Merck). Secondary antibody for SAA was Alexa Flour 488 goat anti-mouse IgG (A-11004, Invitrogen). Iodixanol solution (Optiprep^TM^) was procured from Stemcell Technologies.

### GelMA synthesis and constitution

The GelMA was synthesized through a reaction with methacrylic anhydride based on our previously published work.^[16]^ The resin was constituted in PBS and the photoinitiator components added (separate formulations for each wavelength) as per the concentrations previously shown in Figure 2. For experiments with cells, iodixanol solution was supplemented instead of PBS to be able to match the refractive index range of cells (1.37-1.375) based on existing studies.^[12,20]^ For the tests requiring fluorescence, Rhod-Acr stock in DMSO (stock concentration at 10 mg/ml) was added at 1 µl/ml into the resin formulations, similar to our previous work.^[66]^

### Muscle construct fabrication and culture

C2C12 murine myoblast line was cultured in growth media consisting of DMEM supplemented with 10% w/v FBS and 1% v/v P/S. The cells were passaged at 80% confluency using Trypsin-EDTA and counted via an automated cell counter (LUNA-FX7™, Logos Biosystems). The cells were then resuspended in GelMA resins with LAP at 10^6^ cells/ml (details on optimization of iodixanol concentration are provided in the methods above). The resins were then transferred to 10 mm path length cuvettes and stored at 4°C for 10 min (to allow thermo-reversible gelation of the resin) prior to printing. For printing, the fiber bundle was positioned flush with the cuvette and light projection executed at a light dose increased by 20% to account for light scattering due to cells (optimized from pilot experiments). After printing, the cuvettes were heated at 37°C and the constructs retrieved, washed with PBS to removed the unreacted resin and transferred to 12 well culture plates. The constructs were cultured in 2 ml of growth media (media changes every 48 hours) up to 1 week, followed by differentiation in DMEM containing 2% v/v horse serum and 1% v/v P/S.

### Cell infiltration tests

C2C12 murine myoblasts were encapsulated in GelMA resins with LAP (without Iodixanol) at 10^6^ cells/ml within the 8 well plates (details in the consumables sourcing). 100 µl (∼ 1 mm construct height) of the resin was added to each well and crosslinked within the UV chamber (details in hardware sourcing) with the same light dose as the fiber- crosslinked GelMA. Half of the cell-laden GelMA was then removed using a scalpel and the remaining crosslinked constructs washed with warm PBS (at 37°C) to remove un-initiated LAP. Fresh acellular GelMA (at 37°C) was then added into the empty half of each well plate (i.e., the regions where the cell-laden constructs were removed from), and the well plates stored at 4°C for 15 min. The fiber was used to then crosslink the resins (light doses optimized from Figure S1), and the unreacted resin removed using warm PBS. The constructs were cultured in 150 µl of growth media above the constructs (media changes every 24 hours) up to 96 h, followed by fixation (methods above) and staining for nuclei (DAPI) and F-actin (phalloidin).

### Live/dead analysis and immunohistochemistry

Calcein-AM and propidium iodide were used to stain live and dead cells respectively based on our previous work.^[16]^ For immunostaining, the media was removed and the constructs washed in warm PBS, followed by incubation in 4% v/v PFA for 30 min. The PFA was then removed and the constructs thoroughly washed with PBS. The constructs were then 0.1% v/v Triton-X100 solution for 30 min at room temperature followed by washing thoroughly with PBS. Blocking was then performed in 5% w/v BSA-PBS solution for 1 hour. Primary antibody was diluted in 1% w/v BSA-PBS solution as 1:500 and incubation of the constructs with the primary antibody solution was performed in refrigeration conditions (4 °C) for 12 hours. The constructs were then thoroughly washed with PBS and then incubated for 2 h in secondary antibody solution consisting of 1:500 dilution of the IgG antibodies and 1:1000 for Hoechst and phalloidin. The samples were then thoroughly washed with PBS and imaged in a Confocal microscope (SP8-AOBS-CARS, Leica).

### Shape index calculations

The shape index of the images projected onto the screen was calculated by comparing the size of the actual image to the size of the intended image. For this, the distance (arc length) between the first instance of the minimum gray value and the last instance of the minimum gray value along the arc (red line depicted in Figure 5G) was taken as the actual width of the image. The intended image was superimposed onto the actual image after accounting for the divergence of the light from the fiber bundles (with or without the lenses). The shape index (always < 1) was the ratio of the width of intended image to the width of the actual image.

### Statistical analysis

For studies with three groups, one way Anova with Tukey HSD posthoc tests were used. For studies with two factors, unpaired t-tests (t-tailed) were used for determining the statistical differences.

## Supporting information

FaSt-Light logo projection from fiber bundle with thick cladding

ETH logo projection from fiber bundle with thin cladding

Spoke wheel image projection from the fiber bundle

Projection of window mesh pattern onto rat skin defect

Testing conformation of the mesh on the skin defect

## Supporting information

A supporting information file with Figures S1 to S8 has been provided with the manuscript. Additional videos S1-S3 have also been provided. Data is available at the ETH database: 10.3929/ethz-b-000710181

## Acknowledgements

P.C. acknowledges Spark grant (CRSK-2_220980) and Ambizione grant (PZ00P2_216356) from the Swiss National Science Foundation (SNSF).

## Notes

### Competing Interest Statement

The authors have declared no competing interest.

