## Supplemental info for "Structured light projection using image guide fibers for in situ photo-biofabrication"

Supplemental information consists of Figures S1 to S8

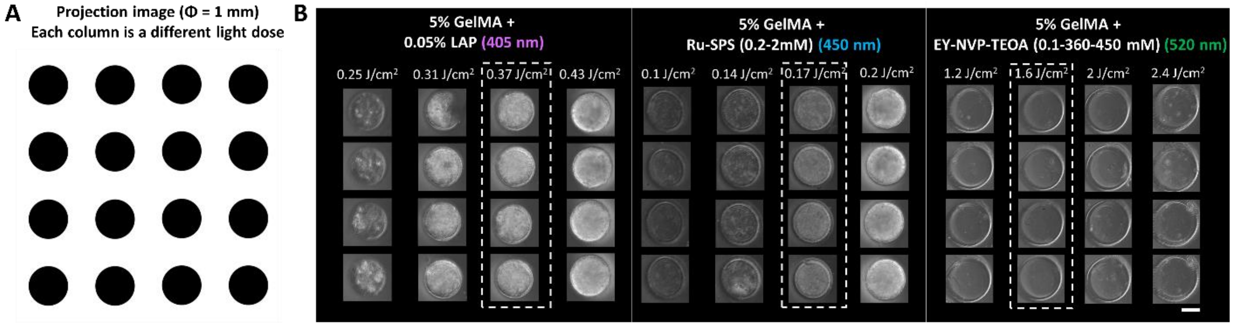


**Figure S1. A.** Projection image for the optimization of the light dose for crosslinking (resins contained within 2 mm path length cuvettes) at each wavelength (photoinitiator components and their components are listed in the images). **B.** Crosslinked images and optimal light doses for each wavelength (marked by dotted rectangles) which result in cylinder diameter closest to the intended design ϕ = 1 mm. Scale bar = 500 µm.

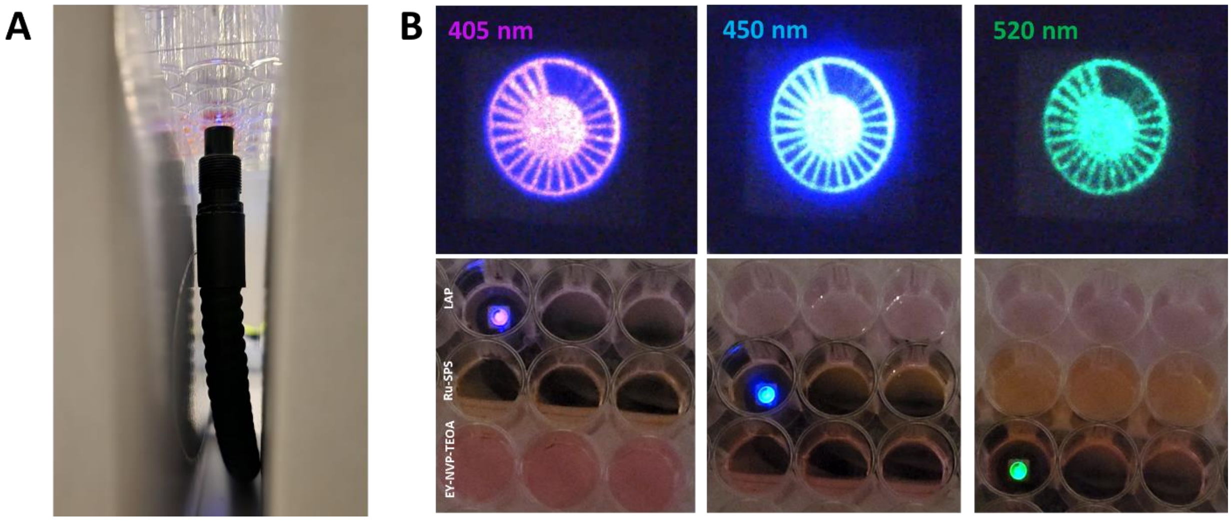


**Figure S2. A.** Setup for the bottom-up projection of images into 48 well plates using the image guide fiber bundles. **B.** Projected spoke wheel patterns at each wavelength within resins (200 µl/well) contained within the well plates.

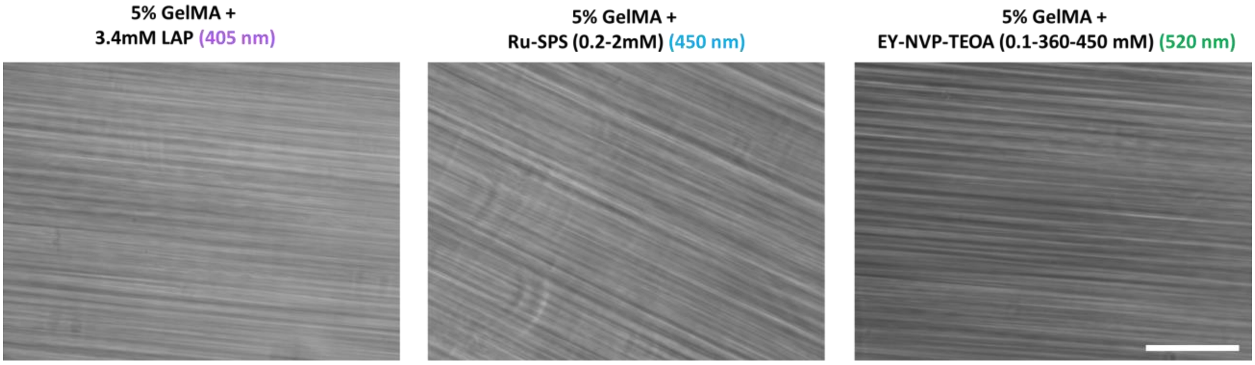


**Figure S3.** Micrographs of microfilament structures within the different photoresin formulations crosslinked using the fiber bundle. Scale bar = 150 μm.

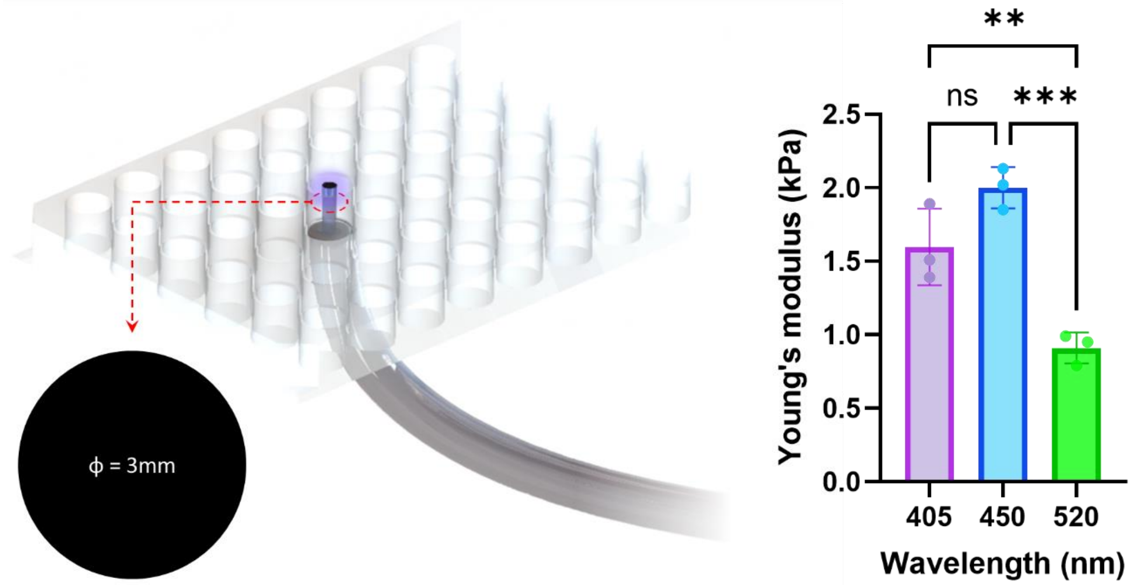


**Figure S4.** Setup for fabrication of cylindrical constructs for mechanical testing (left), where a circular image (diameter ϕ = 3 mm) was projected into well plates containing the photoresin formulations (200 μl/well). ** represent $p<0.01$, *** represent $p<0.001$.

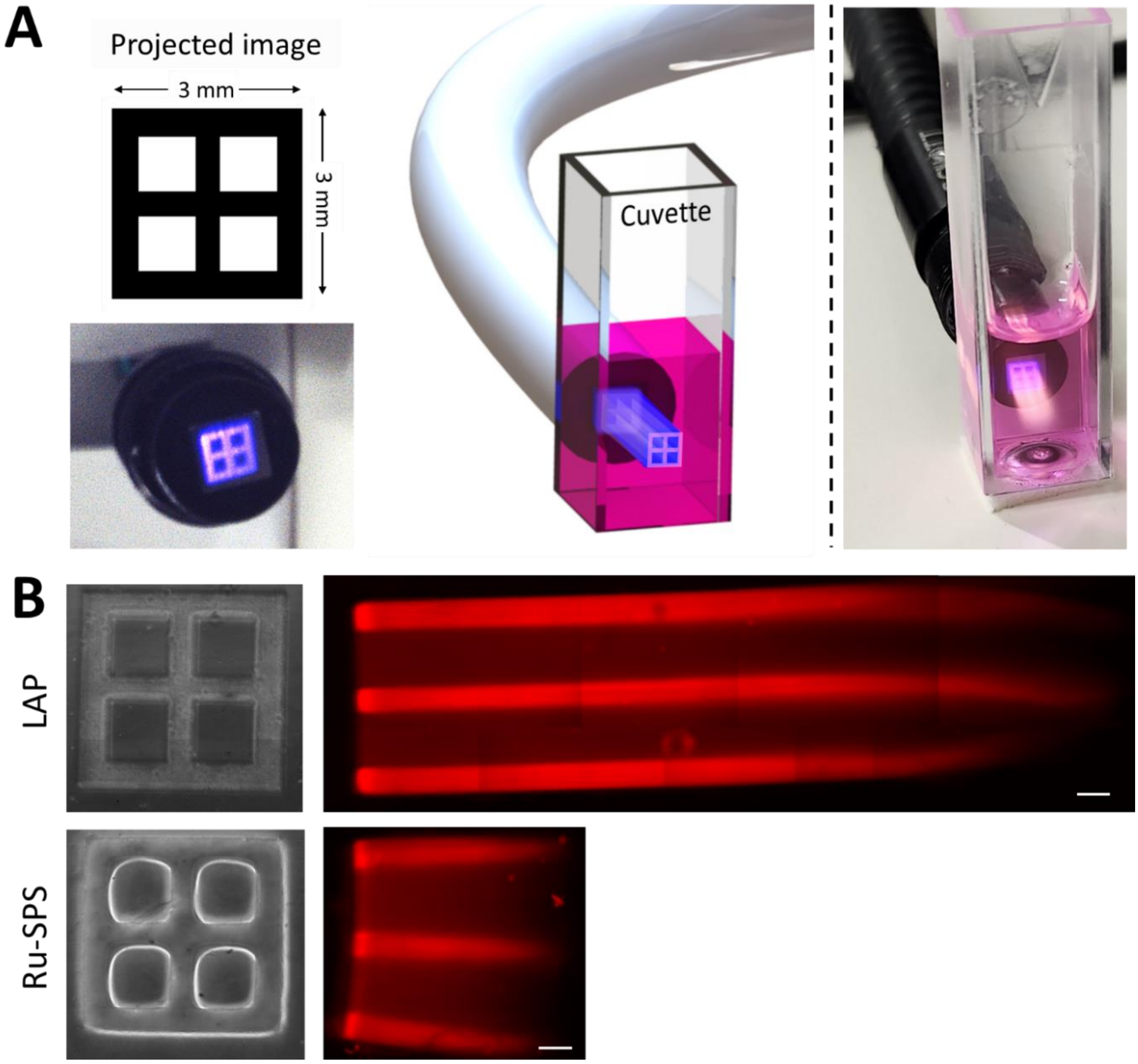


**Figure S5. A.** Projection of image patterns into 10 mm cuvettes. **B.** Sectional views of the crosslinked constructs, which demonstrate higher penetration depth in the LAP-containing photoresins in comparison to those containing Ru-SPS. Scale bar = 500 μm.

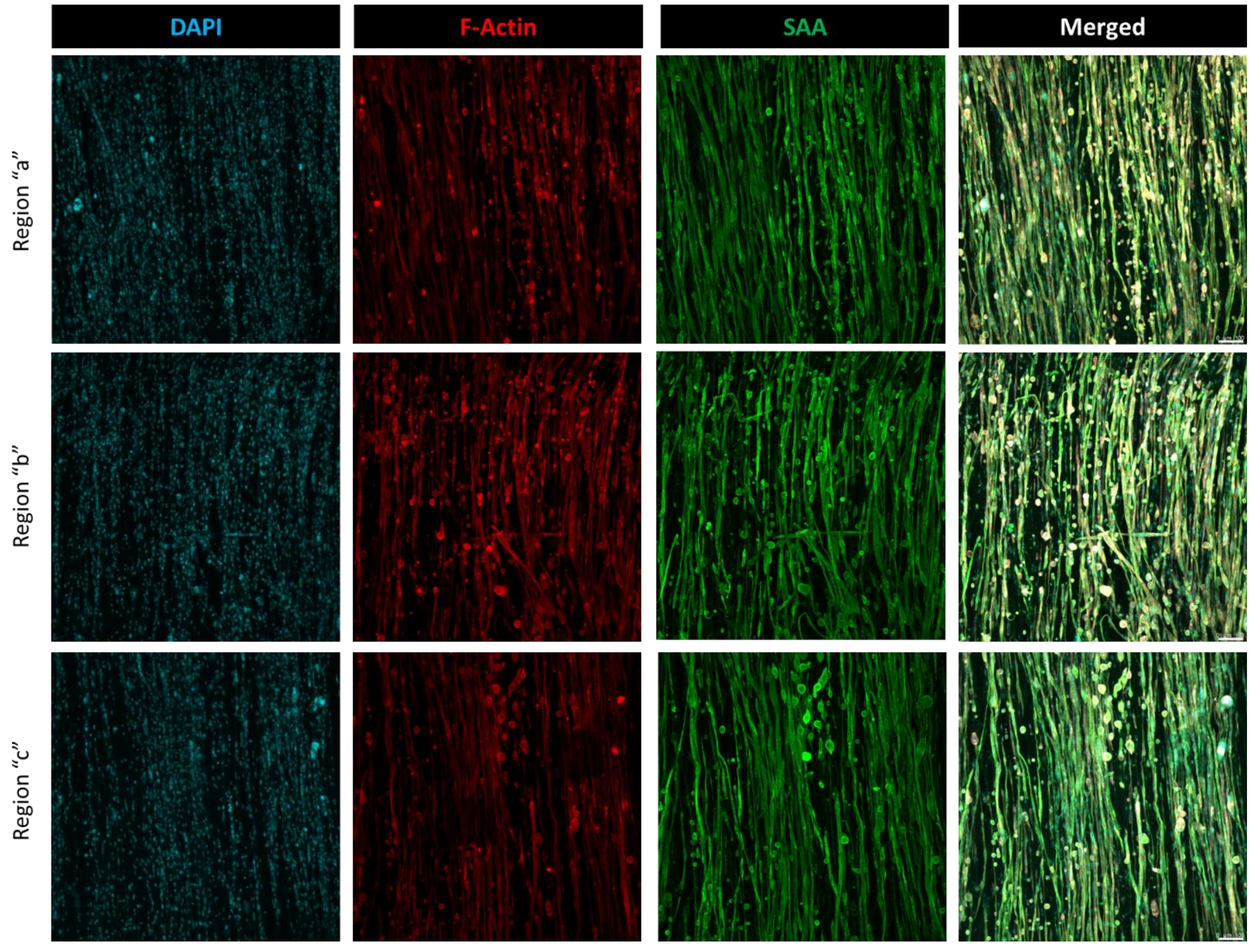


**Figure S6.** Micrographs of myotube distribution in the different regions of the muscle constructs biofabricated using the fiber bundle. All three regions along the constructs demonstrated similar density of myotubes. Scale bar (bottom right in the merged panel) = 100 µm.

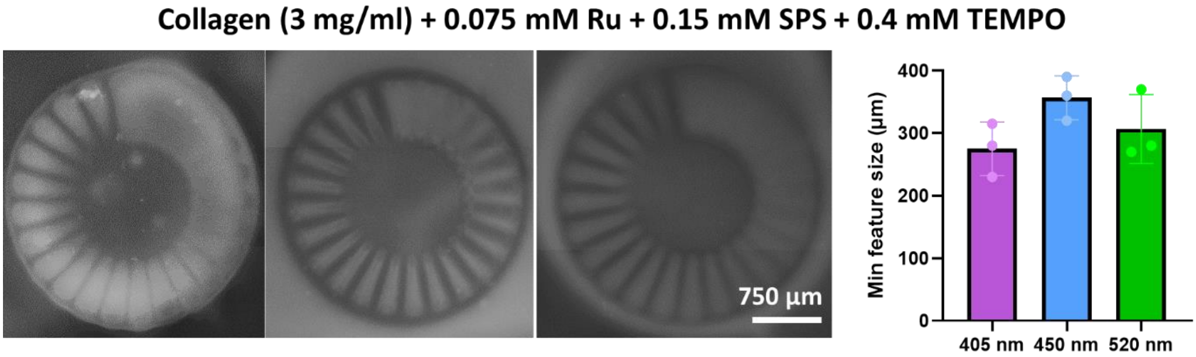


**Figure S7.** Crosslinking of pristine collagen resins containing Ru-SPS photoinitiation system (concentrations listed in the image) using the fiber bundles (projection setup as per Figure S2) and the corresponding minimum feature sizes obtained.
